# Red backgrounds have different effects on electrophysiological responses to fearful faces in groups with low and high autistic tendency

**DOI:** 10.1101/2021.11.21.469184

**Authors:** Eveline Mu, Laila Hugrass, David Crewther

## Abstract

Visual processing differences in the magnocellular pathway have been reported across the autistic spectrum. On the basis that the firing of primate Type IV magnocellular cells is suppressed by diffuse red backgrounds, several groups have used red backgrounds as a means to investigate magnocellular contributions to visual processing in humans. Here, we measured emotional identification accuracy, and compared the P100 and N170 responses from groups with low (*n*=21; AQ<11) and high (*n*=22; AQ>22) Autism Spectrum Quotient (AQ) scores, in response to low (LSF) and high (HSF) spatially filtered fearful and neutral face stimuli presented on red and green backgrounds. For the LSF stimuli, the low AQ group correctly identified fearful expressions more often when presented on a red compared to a green background. The low AQ group also showed red backgrounds reduced the effect of LSF fearful expressions on P100 amplitudes. In contrast, the high AQ group showed that background colour did not significantly alter P100 responses to LSF stimuli. Interestingly, red background reduced the effects of HSF stimuli for the high AQ group. The effects of background color on LSF and HSF facial emotion responses were not evident for the N170 component. Our findings suggest that presenting face stimuli on a red background alters both magnocellular and parvocellular contributions to the P100 waveform, and that these effects differ for groups with low and high autistic tendencies. In addition, a theoretical model for explaining the temporal differences in facial emotion processing for low and high AQ groups is proposed.

## 1. Introduction

Rapid visual processing of fearful facial emotion is important for social functioning and for responding to potential threats in our environments. Facial emotion processing is impaired for individuals with autism spectrum disorder, and these impairments extend to the neurotypical population for groups with high versus low levels of autistic personality traits. Hence it is important to understand the neural basis of rapid facial emotion processing deficits for people with high levels of autistic personality traits. When viewing emotional faces, people with autism (Baron-Cohen et al., 2000; Corbett et al., 2009) and high levels of autistic traits tend to exhibit abnormal activation in amygdala (Nummenmaa et al., 2012), a subcortical region involved in affective processing. One proposal is that abnormalities in the rapidly conducting magnocellular pathways from the retina to the amygdala may underlie facial emotion processing deficits in the broader autistic phenotype (Burt et al., 2017).

Studies on the neural processing of emotional stimuli have implicated the amygdala as a crucial component in mediating such affective processing (Adolphs et al., 2001; Amaral et al., 2003; Batty & Taylor, 2003; Blau et al., 2007; LeDoux, 2003; Morris, 1998; Stein et al., 2014). The amygdala receives input via many visual processing routes, subcortical and cortical, with outputs distributed widely across occipital and parietal regions (Krolak-Salmon et al., 2004; Kveraga et al., 2007; Pessoa, 2010). The most prominent hypothesis regarding the rapid activation of the amygdala is the ‘low road’ hypothesis (LeDoux, 1996), which postulates that the initial analysis of threatening stimuli bypasses the visual cortex utilizing a rapid subcortical pathway projecting from the retina to the amygdala via the superior colliculus and pulvinar (with latencies of ∼80ms) (Garvert et al., 2014; McFadyen et al., 2017; Morris et al., 1999; Tamietto et al., 2012). The alternative ‘high road’ geniculo-V1-amygdala pathway (Pessoa & Adolphs, 2010), demonstrates a latency of ∼140ms (McFadyen et al., 2017; Silverstein & Ingvar, 2015), and recent anatomical research in primate has demonstrated a direct retina-pulvinar-amygdala pathway (Mundinano, 2019). However, there have been few investigations into how amygdala activation modulates cortical visual processing.

The magnocellular and parvocellular pathways work in parallel, processing low-level visual features. The magnocellular pathway is a fast-conducting stream that is insensitive to colour when the luminance is balanced, has higher contrast sensitivity, and is involved in transient attention, while the slower parvocellular pathway performs is colour sensitive, has lower contrast sensitivity, and is involved in sustained responses (Derrington & Lennie, 1984; Kaplan & Shapley, 1986; Laycock et al., 2007; Nassi & Callaway, 2009). It has also been shown that the two early visual pathways are sensitive to different spatial frequencies, with the magnocellular pathway being relatively sensitive to high temporal and low spatial frequency (LSF) information, whilst the parvocellular pathway is more sensitive to low temporal and high spatial frequency (HSF) information. Considering the evidence that fearful stimuli are rapidly processed subcortically (Carr, 2015; Garvert et al., 2014; Johnson, 2005; Morris et al., 1999; Öhman, 2005; Tamietto et al., 2012), it is plausible that this pathway processes coarse, LSF information (Burt et al., 2017). This is consistent with findings that low pass filtered emotional faces activate the amygdala more strongly than do high pass filtered faces (Vuilleumier et al., 2003). This suggests that the magnocellular channels are responsible for driving the rapid salient information to the amygdala. Some researchers (Burt et al., 2017; McCleery et al., 2007) have proposed that less efficient inputs from the magnocellular pathway might contribute to differences in face processing across the autistic personality spectrum.

Event related potentials (ERPs) from EEG recording have superior temporal resolution compared with other brain imaging techniques such as fMRI and are highly effective in identifying the timing of neural responses to emotional information. Two early ERP components studied in affective research are the P100 and N170. The P100 represents a positive early peak approximately 100ms post-stimulus onset. This peak reflects attentional gain (Hillyard & Anllo-Vento, 1998), and seems to be more sensitive to low-level visual information (Mangun, 1995). The N170 component is a negative deflection appearing approximately 170ms post-stimulus. It reflects a structural and featural encoding phase in face processing (Mangun, 1995). The P100 is recorded maximally over lateral occipital-parietal sites (Hillyard & Anllo-Vento, 1998), while N170 originates from regions associated with face and object processing, such as the fusiform gyrus, superior temporal sulcus and inferior, middle and superior temporal gyri (Henson et al., 2003). Visual detection of fearful faces elicits a strong occipital-parietal peak as rapidly as 80ms post-stimulus (Olivares et al. 2015) and 120-140ms post-stimulus (Eimer & Holmes, 2002; Pourtois et al., 2004; Vlamings et al., 2009) compared to neutral faces. This suggests that the amygdala may act as an early alerting mechanism which efficiently redirects visual attention to the threatening stimuli derived from the fast-magnocellular route (Dumas et al., 2013; Ohman, 2005).

Considering magnocellular and parvocellular neurons respond preferentially to LSF and HSF visual input, respectively (Benardete & Kaplan, 1999a, 1999b; Kaplan & Shapley, 1986), it is likely that LSF and HSF filtered faces bias visual processing towards the magnocellular and parvocellular routes, respectively. As such, previous electrophysiological evidence in typically developing populations suggests that P100 amplitudes are greater in response to LSF fearful expressions than to neutral expressions, but HSF emotional expressions do not modulate P100 amplitudes (Pourtois et al., 2005; Vlamings et al., 2009). Some studies have found effects of LSF fearful expressions on N170 amplitudes (Vlamings et al., 2009), while others have not (Holmes et al., 2005; Pourtois et al., 2005). A recent EEG study showed that for participants with low AQ (Autism spectrum Quotient) scores, LSF fearful expressions have larger effects on P100 amplitudes than HSF fearful expressions, yet for people with high AQ scores, fearful expressions tended not to influence P100 amplitudes (Burt et al., 2017). This supports the notion that fearful face processing deficits in ASD could be related to magnocellular pathway abnormalities in processing LSF visual input (Corradi-Dell’Acqua et al., 2014; Laycock et al., 2007).

Single cell studies by Wiesel and Hubel (1966) in lateral geniculate nucleus provided evidence that presenting stimuli on a red background suppresses spiking activity in a class (Type IV) of magnocells (despite the general colour insensitivity of the magnocellular class). These cells have been reported in a number of locations along the magnocellular pathway, including the retinal ganglion cells (de Monasterio, 1978), lateral geniculate nucleus (Wiesel & Hubel, 1966) and striate cortex (Livingstone & Hubel, 1984).

Several studies in human have borrowed from the primate results and have used red surrounds to investigate the effects of suppressing magnocellular firing on human perception and action, and have inferred a similar effect in humans based on behavioural performance change in response to red light, typically a weakened or reduced response in healthy controls (Awasthi et al., 2016; Breitmeyer & Breier, 1994; West et al., 2010; but see Hugrass et al., 2018). West et al. (2010) found a temporal precedence effect for fearful faces that are presented on a green background, but this effect is diminished when the stimuli are presented on a red background of the same luminance. The authors interpreted this finding in terms of suppression of magnocellular input - superior colliculus - pulvinar route to the amygdala. However, it is unlikely that a red surround would disrupt early processing via this route, because unlike Type III magnocellular neurons, Type IV magnocellular neurons do not project to the superior colliculus (de Monasterio, 1978), but project solely to the LGN.

Interestingly, recent studies have shown that red surrounds have different effects on visual processing for groups with atypical magnocellular processing, such as schizophrenia and high trait schizotypy (Bedwell et al., 2013; Bedwell et al., 2006, 2018; Bedwell & Orem, 2008) dyslexia (Chase et al., 2003; Edwards et al., 1996). For example, Bedwell et al. (2013) examined the effects of a red background on P100 responses to a high contrast check pattern. For people with low trait schizotypy, a red background produced the expected reduction in P100 amplitude, whereas for people with high schizotypy, P100 amplitudes did not differ with red and green backgrounds. Similarities in magnocellular functioning in individuals with ASD and schizophrenia (Butler et al., 2005; Kim et al., 2006; Laycock et al., 2007), and the findings that individuals with high AQ and high SPQ scores share a common social factor (Dinsdale et al., 2013; Ford & Crewther, 2014), implies that groups with high versus low AQ groups are likely to show differential processing of stimuli with red versus green backgrounds.

Based on West et al.’s (2010) results, it was predicted that low AQ participants would be less accurate in discriminating between fearful and neutral expressions when LSF face stimuli are presented on a red background, than when presented on a green background. Based on the existing literature (Awasthi et al., 2016; West et al., 2010), it was expected that red background would reduce the effects of LSF fearful expression on P100 and N170 amplitudes in the low AQ group. Finally, combining the evidence that both AQ and schizotypy are associated with magnocellular pathway abnormalities (reviewed Laycock et al., 2007), that the AQ and SPQ scales share a common factor (Dinsdale et al., 2013; Ford & Crewther, 2014) and that red backgrounds have different effects on visual processing for groups with low and high schizotypy (Bedwell et al., 2013), it was anticipated that a red background would not influence the amplitude of early EEG responses to fearful faces in the high AQ group.

## 2. Materials and methods

### 2.1. Participants

Participants were recruited within the university and local community through distributed advertisements and word-of-mouth. The inclusion criteria required participants to be aged between 18 and 40, have normal (or corrected-to-normal) vision, and have no history of neurological conditions. Prior to the EEG session, participants completed an online version of the AQ questionnaire (Baron-Cohen et al., 2001). Of the 135 AQ respondents, 46 individuals who scored either low or high on the AQ questionnaire were recruited to participate in the EEG study at Swinburne University of Technology, Melbourne, Australia. After screening the EEG data for excessive noise (>75μV signals on a high proportion of trials), 43 participants (25 females) were included in the final sample (age range: 18-31 years, *M* = 23.8; *SD* = 4.18). The Swinburne University Human Research Ethics Committee approved the experimental procedures, and all participants provided written, informed consent prior to participation in accordance with the Declaration of Helsinki.

### 2.2. Autism-Spectrum Quotient

The AQ (Baron-Cohen et al., 2001) is a self-report questionnaire measuring the degree to which an adult in the general population with normal intelligence has the traits associated with ASD. The 50-item instrument evaluates social skills, attention switching, attention to detail, communication, and imagination. Allocation into the low and high AQ groups was based on the population mean (*M* = 18.10, *SD*= 10.81) for AQ scores (Ruzich et al., 2015). The low (*n*=21; AQ<11) and high (*n*=22; AQ>22) AQ groups had mean scores of 8.48 (*SD*=2.69) and 27.71 (*SD*=6.17), respectively.

### 2.3. Visual stimuli

Stimuli were created for our 2 (facial emotion) x 2 (spatial frequency) x 2 (background colour) design. The greyscale fearful and neutral face stimuli (see Figure 1) were taken from the 7 different identities (4 female) from the Nimstim Face Set (Tottenham et al., 2009). To create LSF (<2 cycles/degree) and HSF (> 6 cycles/degree) faces, all images were filtered with high-pass and low-pass Gaussian filters (Burt et al., 2017). All pictures were fitted within a frame of 500 × 700 pixels, with the external features (hair, neck and ears) removed. To control for low-level contrast differences between the neutral and fearful faces, only closed-mouth faces were used. The LSF and HSF images were then equated for luminance and RMS contrast using a custom Matlab script (The Mathworks, Natick, MA).

**Figure 1.** Visual stimuli. The task involved the presentation of a scrambled face (1800ms) on a coloured background, followed by a spatially filtered fearful or neutral face (500ms). After the face disappeared, a central fixation cross cued the participant to report the facial emotion. The experiment was separated into four blocks, in which the fearful and neutral face stimuli were a) LSF on a green background, b) HSF on a green background, c) LSF on a red background and d) HSF on a red background.

The tasks were created and displayed using VPixx software (Version 3.20, www.vpixx.com), and presented on a 27 × 48cm LCD monitor (60Hz refresh rate). The luminance of the green background (CIEx = 0.33, CIEy = 0.60) was psychophysically matched to that of the red background (CIEx = 0.33, CIEy =0.60, L = 31.9cd/m^2^).

### 2.4. Equiluminance

The red and green backgrounds were matched for luminance using a centrally presented heterochromatic minimum flicker task, coded in VPixx (VPixx Technologies, Montreal, CA). Red luminance was held constant while subjects adjusted the green luminance to minimize the perception of red-green flicker (Fiorentini et al., 1996). The point of equiluminance was obtained by averaging four adjustment trials. In all subjects, the adjusted green luminance values were close to the physical luminance of the red background (*M difference* =0.54 cd/m^2^, *SE*=0.01 cd/m^2^).

### 2.5. Test procedure

Participants were seated in a quiet dark room, at a viewing distance of 70cm from the screen. All participants started with the flicker photometry task, followed by a short training block of 10 trials prior to the experiment. To prevent fatigue, the experiment was split into four blocks of 120 trials (two blocks each for the red and green background conditions, with LSF and HSF fearful and neutral faces randomised within each block). The order of the blocks was counterbalanced across participants. In total, there were 60 replications for each of the eight experimental conditions (2 background colour x 2 spatial frequency x 2 facial emotion). For all trials a scrambled face (1800ms) was presented before the target face and central fixation cross (500ms). After the face disappeared, participants used a RESPONSEPixx button box, connected to DATAPixx hardware (VPixx Technologies, Montreal, CA), to report whether the expression was fearful (right button) or neutral (left button). The fixation-cross remained on the screen until the participant made a response. Instructions to participants emphasised the importance of accurate decisions over speed. Participants were allowed to rest between blocks. The testing session lasted approximately one hour.

### 2.6. VEP recording and analysis

EEG recordings were made from a 64-channel Quickcap using Scan 4.5 acquisition software (Neuroscan, Compumedics) from parietal, temporal and occipital regions (OZ, O1, O2, O3, P3, P4, P5, P6, P7, P8, PO1, PO2, PO3, PO4, PO5, PO6, PO7 and PO8). Fz was used as the ground electrode site and the linked mastoids were used as the reference channel (Vlamings et al., 2009). EOG electrodes were placed vertically at the upper and lower orbital regions of the right eye to monitor ocular artifact. Electrode impedances were kept below 10KΩ.

EEG data were processed using Brainstorm software (Tadel et al., 2011). Each subject’s data were band-pass filtered from 0.1-30Hz and re-referenced to the average reference. The vertical EOG channels were used to detect and remove eye-blink artifacts from the data using Signal Space Projection. Epochs were extracted from -200 – 400ms relative to stimulus presentation, and baseline subtraction was applied (−200 – 0ms). All epochs containing amplitudes larger than 75μV were removed from the analysis. Separate VEP averages were computed for each of the 2 (spatial frequency) by 2 (background colour) by 2 (facial emotion) conditions, for the low and high AQ groups. On average, 57.6 (*SD* = 3.4) out of 60 trials in each condition survived data cleaning processes, with no significant differences in the number of trials retained across conditions (all comparisons *p*>0.32), or across participants in the low versus high AQ groups (all comparisons *p*> 0.16).

To improve the signal with respect to noise, the mean responses were extracted for a cluster of occipitotemporal electrodes from the right hemisphere (P8, PO8). These electrodes produced the greatest amplitude P100-N170 responses, and the electrode locations are consistent with previous literature showing a right hemisphere advantage for the processing of faces and emotion expressions (Burt et al., 2017; Bruno Rossion, 2014; Vlamings et al., 2009). P100 and N170 amplitudes were then extracted using a routine programmed in LabVIEW (National Instruments, USA). P100 amplitude was defined as the maximum amplitude in the time-window from 80 to 140ms after stimulus presentation. N170 amplitude was defined as the negative peak amplitude in the time-window from 150 to 210ms after stimulus presentation. The data were screened for outliers. The data were screened for outliers. For the analysis of P100 amplitude, data from one low AQ participant was excluded due to an outlier for the red background LSF neutral condition.

The peak amplitude and latency data were analysed using SPSS Statistics software (SPSS, Version 20, IBM) and JMP (Version 10.0). In order to investigate the effects of background (green, red) on the ERP emotional responses (fear, neutral) with respect to AQ scores (low, high), we conducted separate 2 (background) by 2 (emotion) by 2 (AQ) mixed design ANOVAs for LSF and HSF stimuli, with P100 and N170 amplitudes as the dependent variables. Separate analyses were conducted for LSF and HSF stimuli, on the basis that this reasonably separates magnocellular and parvocellular contributions (Pourtois et al., 2005; Vlamings et al., 2009). Bonferroni corrections (*α* = 0.0125) for multiple comparisons were applied to all follow-up t-tests.

## 3. Results

### 3.1. Behavioral results

Means and standard errors for emotion identification accuracies are presented in Figure 2. Accuracy was defined as the percentage of trials when fearful and neutral facial expressions were correctly reported. For both AQ groups, mean accuracy was above 90% for all conditions. Response times were not subject to statistical analysis as participants were instructed to decide as accurately as possible, only after the target face had disappeared (in order to minimise movement artifact during EEG recording). Thus, the values obtained would not truly reflect reaction times to the stimuli. Due to corrupted and/or missing key-press files, only data from 20 low AQ and 20 high AQ participants were included in the analyses.

**Figure 2.**
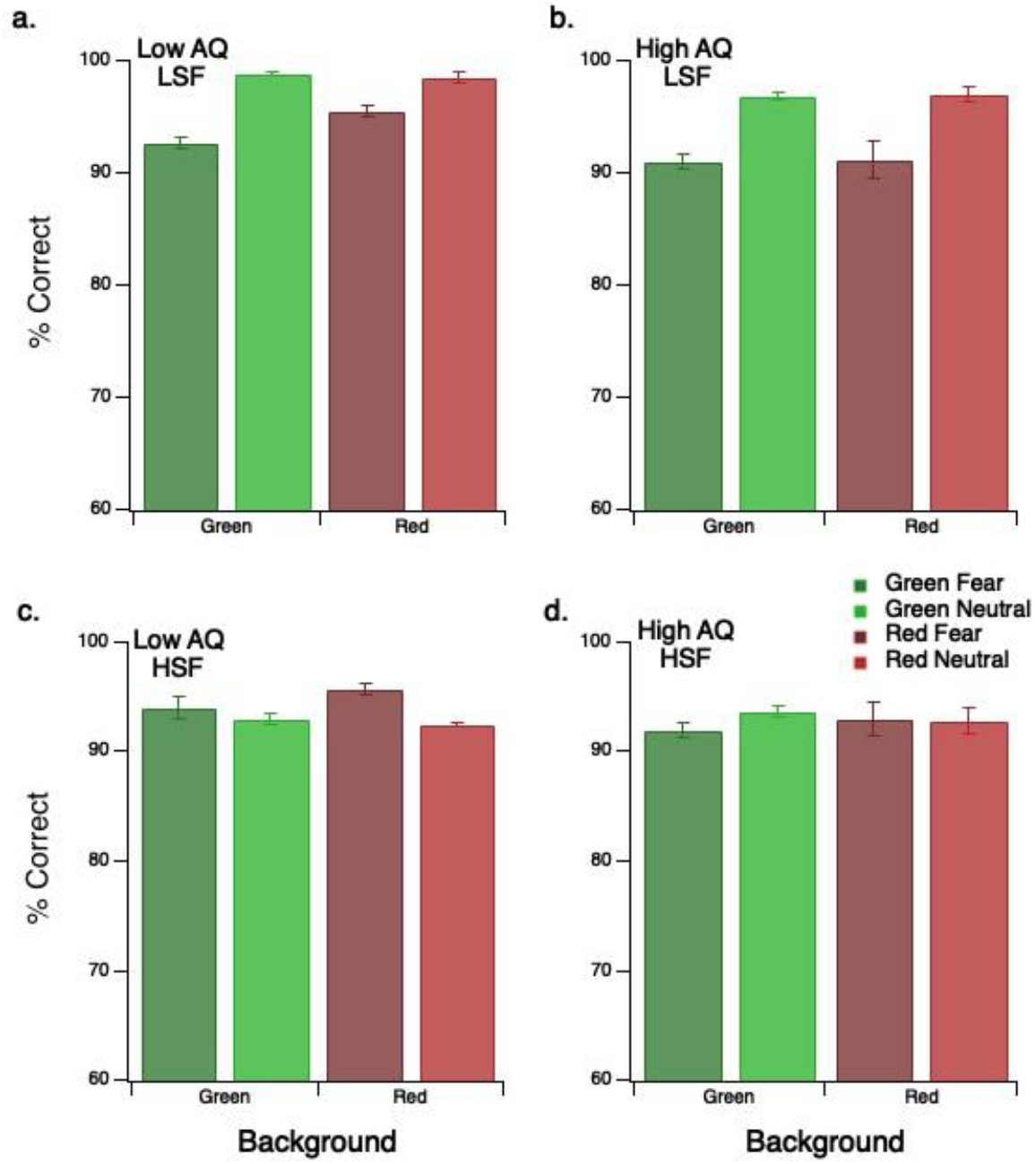
Mean accuracy levels for emotion identification. Results are presented in separate panels for the low AQ group with (a) LSF and (c) HSF stimuli on green and red backgrounds, and for the high AQ group with (b) LSF and (d) HSF stimuli on green and red backgrounds. Results for the fearful and neutral faces are presented in the darker and lighter bars, respectively. Error bars represent within-subject 1 SEM.

For LSF conditions, the three-way (background x emotion x AQ) ANOVA produced a significant main effect of emotion (*F*(1,37)=12.45, *p*=0.001, 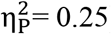), with greater accuracy for neutral compared to fearful faces. There was also a significant interaction between background, emotion and AQ (*F*(1,37)=4.45, *p*=0.041, 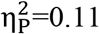). To explore this interaction, two separate 2 (background) x 2 (emotion) ANOVAs were performed for the low and high AQ groups. For the LSF condition, there was a significant background and emotion interaction for the low AQ group (*F*(1,19)=10.41, *p*=0.004, 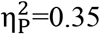), with greater accuracy for fearful faces presented on a red background than on a green background. For the high AQ group, there was a significant main effect of emotion (*F*(1,18)=6.87, *p*=0.017, 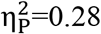), with greater accuracy for LSF neutral faces across both background conditions.

For the HSF conditions, no significant main effects or interactions were evident for either the low or high AQ group.

### 3.2. VEP results

Grand average ERPs for fearful and neutral, LSF and HSF faces are presented in Figure 3, with separate panels for LSF and HSF conditions for low AQ (Figure 3a and 3c) and for high AQ (Figure 3b and 3d) groups.

**Figure 3.**
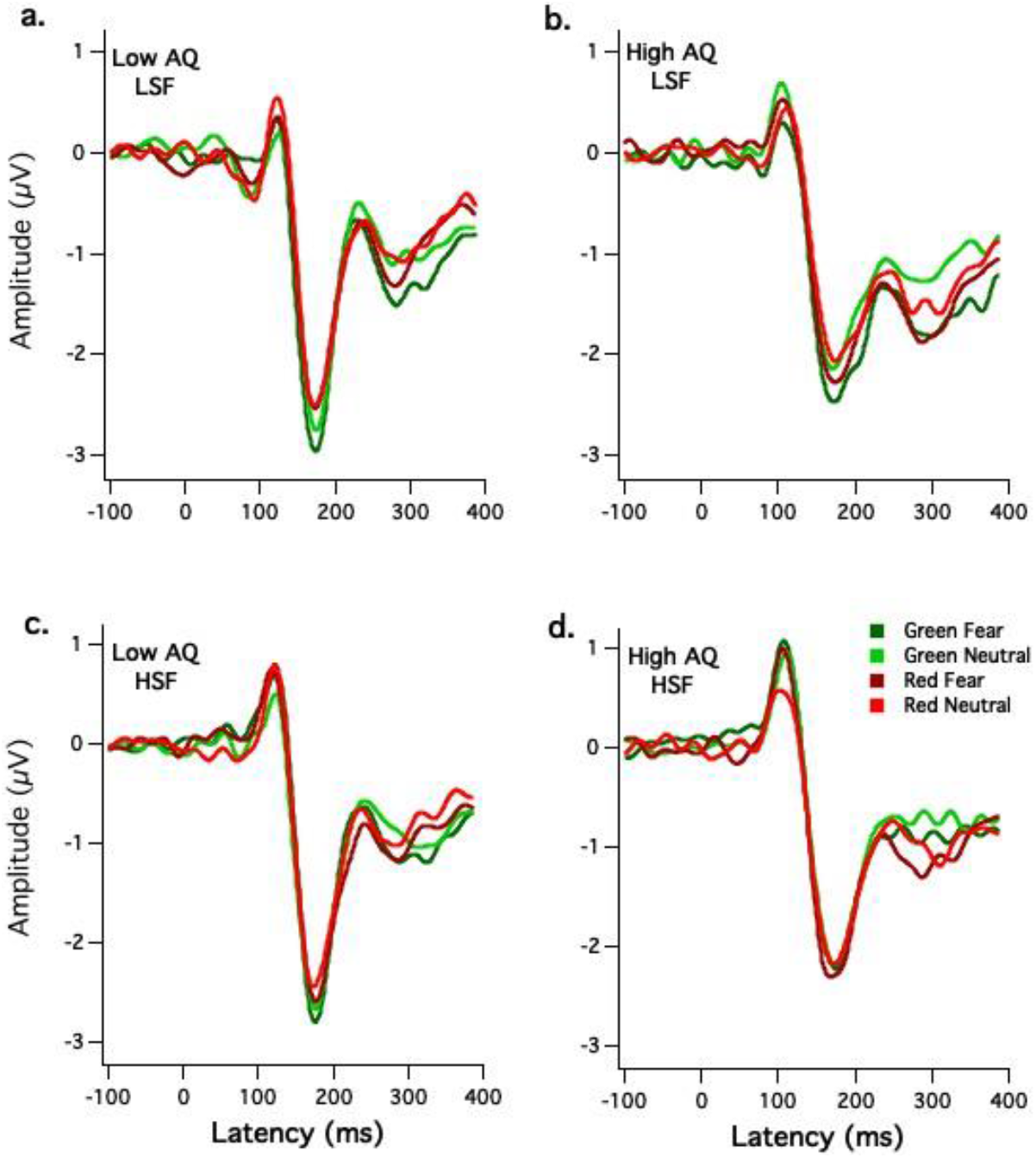
Grand averaged cluster waveforms in response to LSF and HSF fearful and neutral faces. The separate panels illustrate responses from the low AQ group for (a) LSF and (c) HSF stimuli, and the high AQ group for (b) LSF and (d) HSF stimuli. For the red background conditions, the dark and light red lines represent fearful and neutral faces, respectively. For the green background conditions, the dark and light green lines represent fearful and neutral faces, respectively.

#### 3.2.1. P100 amplitude

Figure 4 illustrates the effects of background colour on P100 amplitudes for the low and high AQ groups with LSF (Figures 4a and 4b) and HSF (Figure 4c and 4d) face stimuli. Separate 2 (background) x 2 (emotion) x 2 (AQ) ANOVAs were performed for the LSF and HSF conditions.

**Figure 4.**
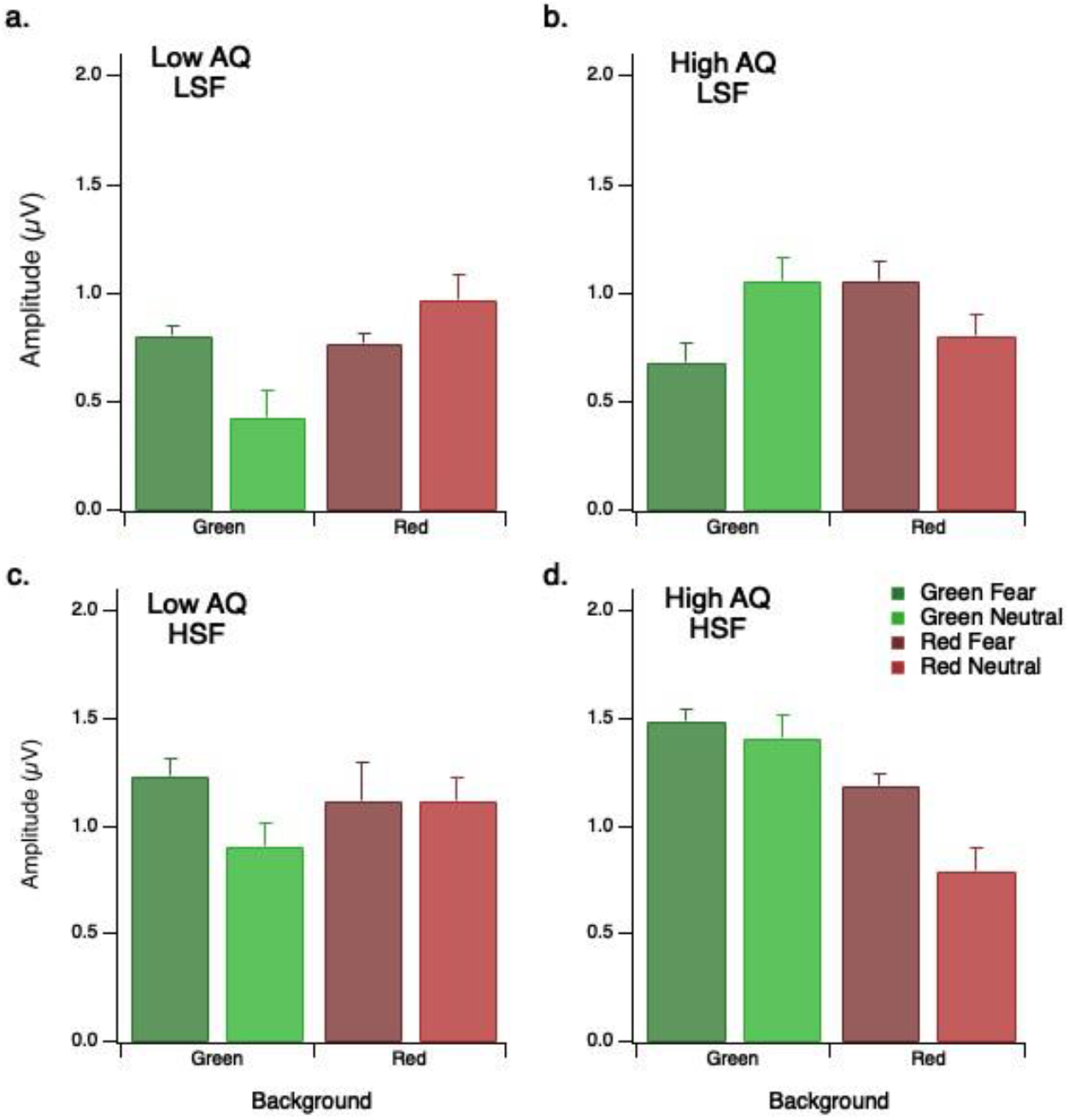
Mean P100 amplitudes. Results are presented in separate panels for the low AQ group with (a) LSF and (c) HSF stimuli, and for the high AQ group with (b) LSF and (d) HSF stimuli. Results for the fearful and neutral faces are presented in the darker and lighter bars, respectively. Error bars represent within-subject 1SEM.

For the LSF condition, there was a significant three-way interaction between the effects of background colour, emotion and AQ *F*(1,40)=4.29, *p*=0.045, 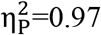). To investigate this interaction, separate follow-up tests of simple main effects were conducted for the low and high AQ groups (Bonferroni corrected). For the low AQ group, mean P100 amplitudes were higher for LSF fearful faces than LSF neutral faces when the background was green 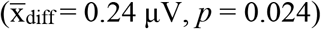, but not when the background was red 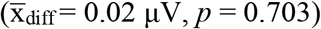. For the high AQ group, mean P100 amplitudes for LSF fearful and neutral faces differed in the opposite direction, but not significantly, for either the green background 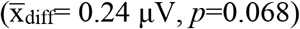 or red background 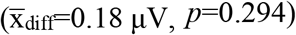 conditions.

For the HSF condition, the main effect of emotion was approaching significance (*F*(1,41)=3.86, *p*=0.056), with a tendency for P100 amplitudes to be greater for fearful faces compared to neutral faces. The interaction between background and AQ was also approaching significance (*F*(1,41)=3.51, *p*=0.068). To explore this interaction, follow-up tests of simple main effects were conducted (Bonferroni corrected). On average, P100 amplitudes were higher in response to stimuli presented on a green background than on a red background for the high AQ group 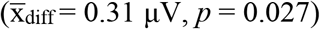 but not for the low AQ group 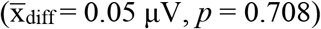.

##### 3.2.1.1. Principle component and factor analysis

To obtain a clearer understanding of the relationships between the stimulus variables as they apply to the hypothetical framework, the P100 amplitudes for LSF fearful and neutral stimuli presented on green and red backgrounds were subjected to a post-hoc principal components analysis (PCA) using maximum likelihood estimation/Varimax rotation. Due to the well-reported relationship of LSF and magnocellular functioning, and the aim of the current study, we conducted this further analysis on LSF stimuli only. Components with Eigenvalues >1.0 were retained as substantial representations of the variation in the model. Component loadings below 0.3 were suppressed in order to report only important component contributions (Tabachnick and Fidell, 2013).

Initially, the AQ scores were included in the PCA in order to establish which other variables that the AQ scores related to. The rotated component loading for P100 amplitudes to LSF stimuli plus AQ are presented in Table 1 below. Scores below 0.3 are not shown. The PCA resulted in a three-component solution (explaining 77% variance), with only component 1 being AQ dependent.

**Table 1.** Rotated component loadings for LSF P100 amplitudes using Maximum Likelihood/Varimax methods

|  | Component 1 | Component 2 | Component 3 |
| --- | --- | --- | --- |
| AQ score | 0.500 |  |  |
| Green LSF Fear |  | 0.788 |  |
| Green LSF Neutral | 0.801 |  |  |
| Red LSF Fear | 0.322 |  | 0.548 |
| Red LSF Neutral |  |  | 0.557 |

Note: Scores below 0.3 are not shown in order to report only important component contributions.

The individual component scores were calculated and presented as a 3D scatter plot (see Figure 5). Membership of the low AQ and high AQ groups is shown by grey cubes and black spheres respectively. The separation of most of the low AQ data points from the high AQ data points in the 3-factor space clearly shows that autistic tendency plays a role in the strength of the P100 response for LSF stimuli.

**Figure 5.**
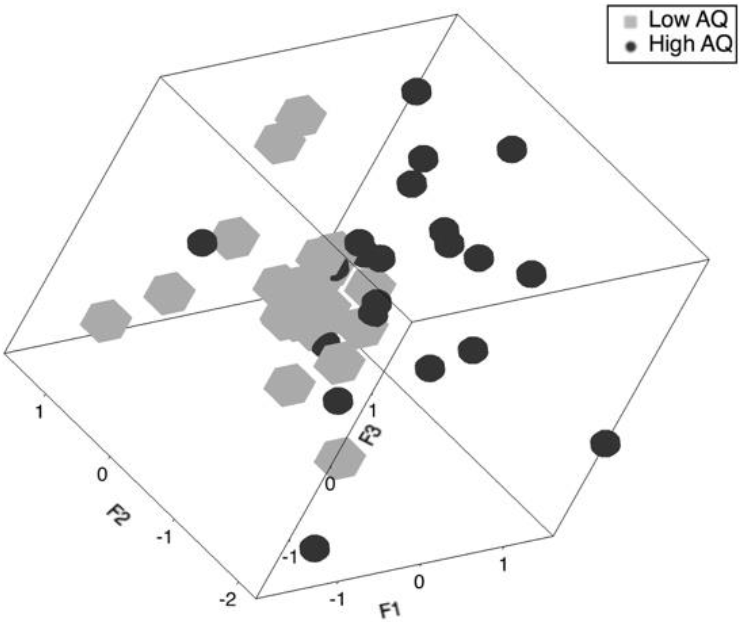
3D component plot of low AQ (grey cube) and high AQ (black sphere) groups plotted as a function of the 3 components. The view chosen shows a clear separation of low and high AQ data points

**Figure 6.**
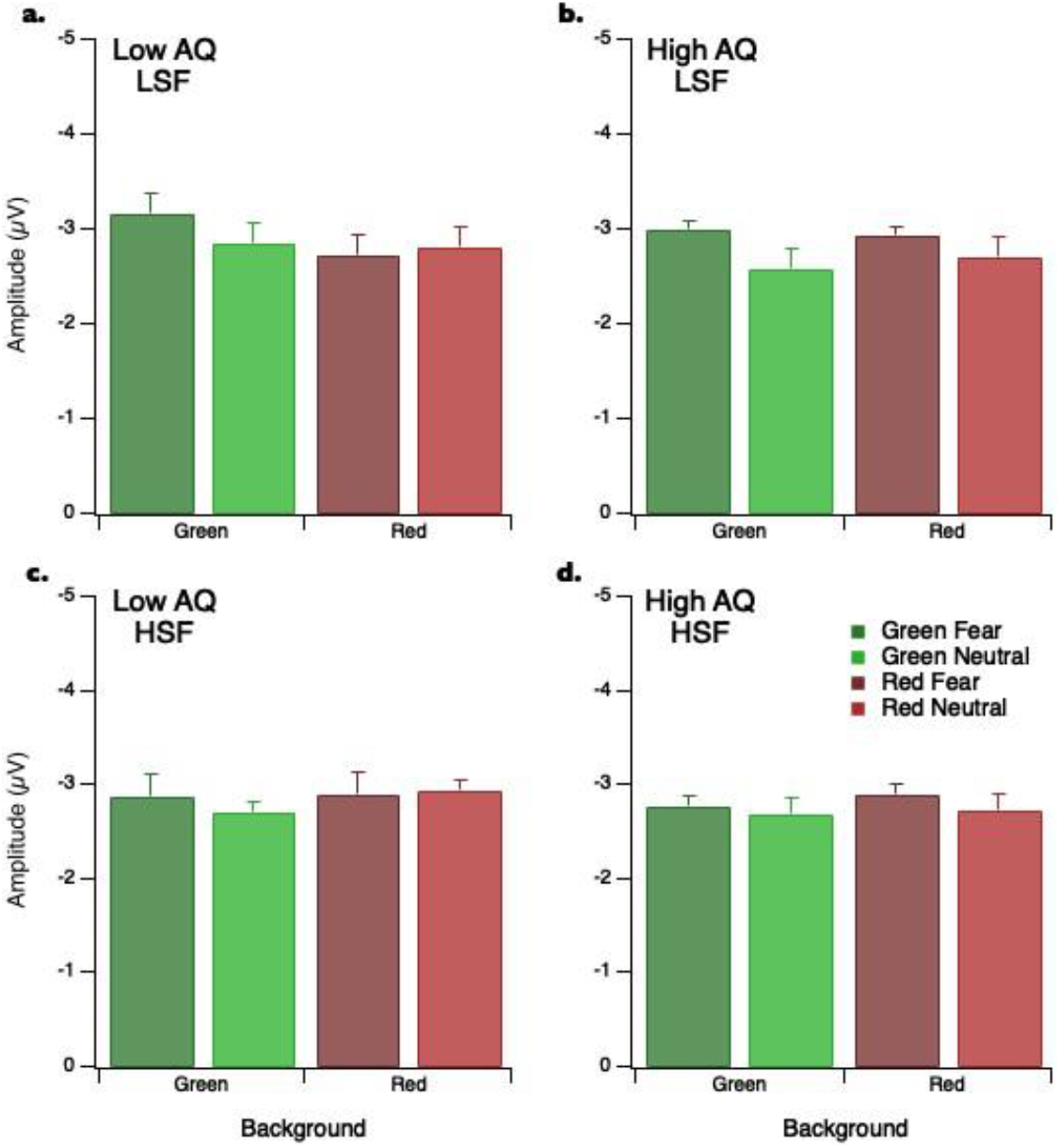
Mean N170 amplitudes. Results are presented in separate panels for the low AQ group with (a) LSF and (c) HSF stimuli, and for the high AQ group with (b) LSF and (d) HSF stimuli. Results for the fearful and neutral faces are presented in the dark and light grey bars, respectively. Error bars represent within-subject 1SEM.

To further explore factors underlying AQ differences, factor analysis was performed separately for low and high AQ groups, as shown in Table 2 below. Principal components were used as the factoring method with prior communality indexed by squared multiple correlations (SMC), and rotation using Varimax. Only two components reached significance on the Kaiser criterion and hence a two-component solution was developed. For the low AQ group, the first factor coded for fearful faces across red and green backgrounds, while the second factor coded neutral faces across red and green backgrounds (with negative correlations between the amplitudes). In contrast, for the high AQ group, the first factor coded predominately for green backgrounds, with a weaker input from the red neutral variable while the second factor coded for red backgrounds, across all facial emotional expressions.

**Table 2.**
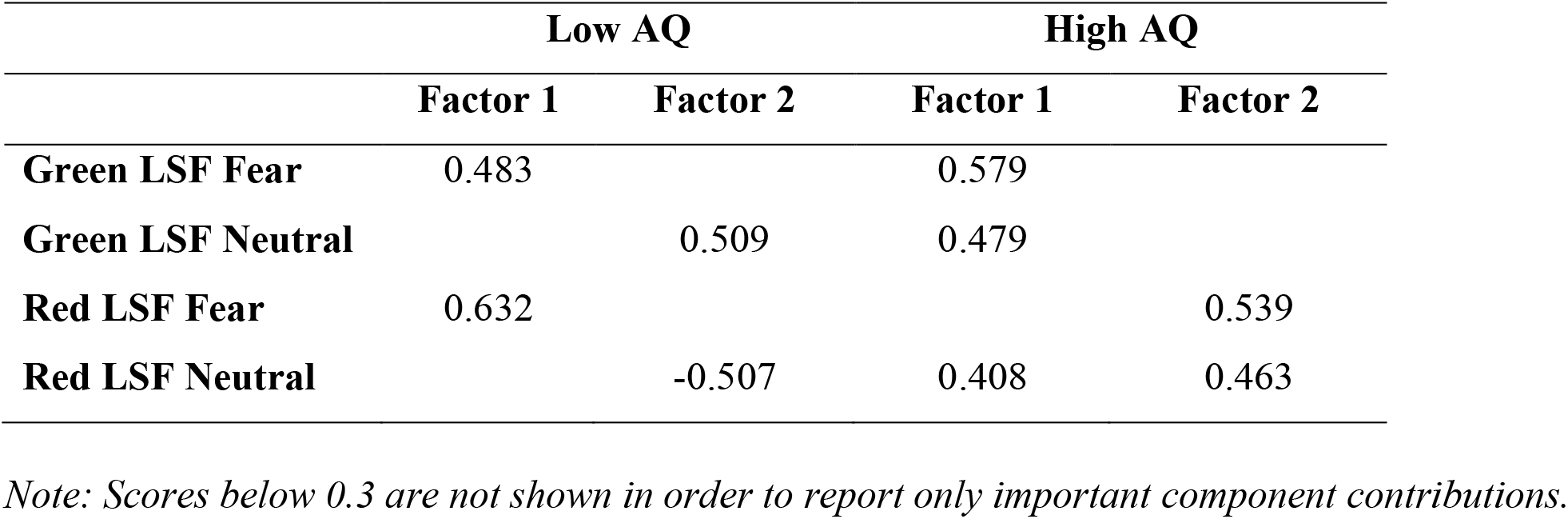
Separate rotated exploratory factor loadings with Varimax rotation for low and high AQ groups

|  | Low AQ |  | High AQ |  |
| --- | --- | --- | --- | --- |
|  | Factor 1 | Factor 2 | Factor 1 | Factor 2 |
| Green LSF Fear | 0.483 |  | 0.579 |  |
| Green LSF Neutral |  | 0.509 | 0.479 |  |
| Red LSF Fear | 0.632 |  |  | 0.539 |
| Red LSF Neutral |  | -0.507 | 0.408 | 0.463 |

*Note: Scores below 0.3 are not shown in order to report only important component contributions.*

In summary, the analysis demonstrated that the strength of P100 responses for LSF face stimuli is dependent on autistic tendency. It also revealed a different factor structure in which for the low AQ group the factors separated on the basis of emotion while for the high AQ group, they separated more on the basis of background colour.

#### 3.2.2. N170 Amplitude

Bar graphs for mean N170 amplitudes, separated by autistic tendency group and spatial frequency are presented in Figure 5. For the LSF condition, the 2 (background) x 2 (emotion) x 2 (AQ) ANOVA produced a significant main effect of emotion (*F*(1,41)=8.201, *p*=0.007, 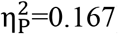), with greater N170 amplitude for fearful faces compared to neutral faces. There were no other main effects or interactions.

For the HSF condition, there were no significant main effects or interactions.

## 4. Discussion

This is the first electrophysiological investigation into the effects of a red background on early evoked activity to fearful and neutral facial expressions for groups with low and high autistic tendency. For the LSF face stimuli, there was a complex interaction between the effects of background colour, facial emotion and autistic tendency on P100 amplitudes. This suggests that the effects of a red background on early evoked responses to fearful versus neutral faces may vary for groups with low and high levels of autistic tendency. These effects of background color on responses to facial emotion were not evident for the N170 component. Hence, our discussion focuses on interpreting the P100 responses and providing a theoretical model for explaining the differences in facial emotion processing for groups with low and high AQ scores.

The behavioral data suggest that both the low and high AQ groups were able to accurately identify the facial expressions as neutral or fearful, regardless of the background colour or spatial frequency of the stimuli. For the low AQ group, we found that LSF fearful expressions were identified correctly more often when the faces were presented on a red background than on a green background. This was initially a surprising result in light of West et al.’s (2010) finding that a red background extinguishes the temporal precedence for detecting fearful facial expressions. However, given the length of the stimulus exposure duration (500ms), our behavioral results are unlikely to reflect rapid processing mechanisms that are important for temporal order judgements.

Previous studies have suggested that the magnocellular pathway allows for rapid transmission of salient information, such as threat, to the amygdala (de Gelder et al., 2011; Méndez-Bértolo et al., 2016; Morris et al., 2001; Vuilleumier et al., 2003). On the basis that magnocellular responses are biased towards LSF input (Benardete & Kaplan, 1999a, 1999b; Kaplan & Shapley, 1986), and that type IV magnocellular cells are suppressed by red backgrounds (Wiesel & Hubel, 1966), we used both spatial frequency and background colour to probe magnocellular involvement in P100 responses to fearful faces. For the low AQ group, the finding that P100 amplitudes were greater for LSF fearful versus LSF neutral faces on a green background supports evidence of a magnocellular pathway involvement in the rapid processing of threat relevant stimuli. Furthermore, our finding that this difference is extinguished when the faces are presented on a red background indicated that suppressing magnocellular input (specifically Type IV magnocellular input) reduces early processing for threatening versus neutral face stimuli. However, these effects were relatively small, which indicates that using a red background does not greatly influence the variance in P100 responses to emotional faces. By contrast, for the high AQ group, there was no significant difference in mean P100 amplitudes for LSF fearful and LSF neutral faces, regardless of whether they were presented on green or red backgrounds. This indicates that for those with high levels of autistic tendency, Type IV magnocellular cells are unlikely to contribute to early processing differences for fearful versus neutral faces.

There are similar magnocellular abnormalities in both AQ and schizotypy (reviewed Laycock et al., 2007), and the AQ and SPQ scales share a common factor (Dinsdale et al., 2013; Ford & Crewther, 2014). Our finding that a red background had different effects on P100 responses to LSF faces for groups with low and high AQ appears to parallel those of Bedwell et al. (2013) who found a red background to reduce P100 amplitude for people with low schizotypy, but no change in P100 amplitude to the red background for people with high schizotypy. Findings from the current study and Bedwell et al. imply that a red background may have different effects on individuals with differing magnocellular function, such as those with low and high AQ. However, such comparisons should be considered with caution as the stimuli employed in the current study and Bedwell et al. are not identical. Specifically, considering we presented facial emotional stimuli we can assume that subcortical projections to the amygdala may influence P100 amplitude, whereas Bedwell et al. presented checkerboard stimuli, which are unlikely to drive a strong amygdala response.

There are also recent reports in primate of wide-field retinal ganglion cells that project directly to the medial sub-division of the inferior pulvinar (Kwan et al., 2019), findings yet to be confirmed in human. For instance, if input to the amygdala is dominated by the LGN-cortical pathways, one might expect a red background to suppress rapid amygdala reactivity to affective stimuli. While there is certainly mixing of information from magnocellular and parvocellular pathways after V1, and there is a strong V1 – pulvinar – extrastriate cortex feed of information (Ahmadlou et al., 2018; Lakatos et al., 2016), such circuitous associations would be arguably less spatial frequency specific and slower (McFadyen et al., 2017), thus less likely to influence P100 responses to emotional face stimuli. In comparison, direct pulvinar–hMT connections would be expected to involve rapid processing (Kwan et al., 2019). If input to the amygdala is strongly influenced by these subcortical pathways, one would not expect red backgrounds to reduce rapid amygdala reactivity to fearful facial expressions.

Interestingly, however, the PCA/factor analysis performed on the P100 responses for LSF face stimuli showed a factor structure separated on the basis of stimulus emotion for the low AQ group, but on the basis of background color for the high AQ group. This indicates that we cannot exclude the notion that Type IV magnocellular cells are likely to contribute to early processing differences for those with high levels of autistic tendency. Rather, we need to examine how magnocellular inputs to the dorsal cortical stream (the source of the P100 response) might differ anatomically or physiologically as a function of autistic tendency.

A potential mechanism to explain general differences in P100 amplitude responses for the low and high AQ group to facial emotional stimuli is an amygdala-driven contrast-gain modulation. Contrast gain effects of amygdala hyper-response to fearful expressions in humans with autism (Amaral et al., 2003), and in animal models of ASD (Markram et al., 2008) are reported to increase hMT and extrastriate early response. Tadayonnejad et al (2016) used effective connectivity analysis of the pulvinar to show that people with generalised social anxiety disorder demonstrated causally increased influential dynamics between pulvinar and higher order visual cortical regions. Similarly, pulvinar manipulation of occipital cortex visual responses is supported by studies on pharmacological agonism and antagonism of pulvinar (Purushothaman et al., 2012) and also the model of amygdala modulation of visual response via pulvinar and thalamic reticular (TRN) (John et al., 2016). Furthermore, such modulation of thalamic gain by amygdala stimulation has received a boost through optogenetic studies in mice (Aizenberg et al., 2019), where optogenetic activation of amygdala amplified tone evoked responses in auditory cortex.

Thus, a model consistent with red backgrounds producing different effects on P100 amplitudes for high versus low AQ individuals can be plausibly constructed via a combination of 1) a hyper responsive amydgala reaction to fear or anxiety generating situations in those with ASD or high AQ (Dalton et al., 2005; Markram et al., 2008); 2) response gain modulation of pulvinar by direct amydgala projections to the TRN; 3) red background suppression of LGN thalamic drive to hMT causing a greater balance of pulvinar-hMT drive and a resultant fear driven emotional attention. Therefore, the differential effects seen with high versus low AQ groups may be explainable through a different weighting of thalamic inputs to parietal cortex through pulvinar and LGN, or from a differential degree of amygdala driven contrast gain modulation, exerted at the pulvinar, for those with high versus low AQ scores.

The literature focuses on the idea that the suppressive effects of red backgrounds are only related to magnocellular functioning (Awasthi et al., 2016; Breitmeyer & Breier, 1994; West et al., 2010; Wiesel & Hubel, 1966), while not considering if there are effects on the spatially and chromatically sensitive parvocellular system. To this point, we found that a red background produced a reduction in P100 amplitudes with HSF stimuli for the high AQ group. This finding is reminiscent of the results of Hugrass et al. (2018), who examined the effects of red background on magnocellular and parvocellular non-linear VEP signatures and found red background to suppress parvocellular generated temporal nonlinearity VEPs, and not did influence magnocellular generated VEP signatures. Considering the parvocellular pathway is highly sensitive to (red/green) color, Hugrass et al. suggested that red backgrounds may increase temporal sensitivity in the parvocellular pathway, with an immediate prediction of enhanced L-M color fusion frequencies. Taken together, one should be cautious when utilising red backgrounds to study magnocellular functioning as they may affect parvocellular functioning too.

In conclusion, we compared ERPs in response to fearful and neutral faces for groups with low and high levels of autistic tendency. We used red and green backgrounds and LSF and HSF stimuli in order to probe the role of magnocellular visual input in driving early cortical responses to fearful stimuli. For the low AQ group, P100 amplitudes were higher for the LSF fearful than neutral face when the stimuli were presented on a green background, but not when the faces were presented on a red background. However, for the high AQ group, P100 amplitudes were higher for HSF fearful than neutral faces when presented on a green background, but not when faces were presented on a red background. Our findings suggest that presenting face stimuli on a red background alters both magnocellular and parvocellular contributions to the P100 waveform, and that these effects vary for groups with low and high levels of autistic tendency

## 5. Author contribution statement

EM created the experimental design, performed testing and data collection, and drafted the manuscript. EM and LH analysed the data. LH and DC contributed to manuscript editing. All authors contributed equally to interpreting the results.

## 6. Conflict of interest statement

The authors declare that research was conducted in the absence of any commercial or financial relationships that could be construed as a potential conflict of interest.

## Notes

### Competing Interest Statement

The authors have declared no competing interest.

